# The use of X-ray Computed Tomography revolutionises soil pathogen detection

**DOI:** 10.1101/2024.11.18.624060

**Authors:** Eric C. Pereira, Christopher Bell, Peter E. Urwin, Saoirse Tracy

## Abstract

Potato cyst nematodes (PCN), namely *Globodera pallida*, pose a significant threat to potato production worldwide. Accurately quantifying nematode population densities is crucial for effective management strategies. However, traditional detection methods are time-consuming and often imprecise. X-ray computed tomography (CT) offers high-resolution, non-destructive and three-dimensional (3D) imaging frequently used to visualise internal structures of objects, materials or biological tissues, allowing for detailed analysis of their composition, defects, and morphology. Here, we demonstrate the effectiveness of X-ray CT in detecting and quantifying PCN cysts in different soil types. The ability to achieve 3D imaging and volume quantification of PCN cysts from soils allows accurate enumeration of their egg content. To explore future potential, we propose that when integrated with AI-led quantification of X-ray CT-generated cyst counts, this method could evolve into a next-generation technology to accurately and rapidly quantify these damaging pests in agricultural soils. Combined with the potential to co-analyse other organisms within the same sample, we propose X-ray CT as a fundamental tool for comprehensive soil health assessments and sustainable pest management in agriculture.

**Author Summary:** Potato cyst nematodes are a major threat causing significant crop losses worldwide. Traditionally, detecting this pathogen in soil is challenging, often requiring time-consuming and imprecise. Our study explores a new approach using X-ray computed tomography (CT), a powerful imaging tool that creates detailed 3D images *in situ*. We observed that X-ray CT can effectively detect and measure the cysts nematodes, even within different soil types. This detailed analysis allows us to determine the cyst structures and estimate the eggs contained. In the future, combining X-ray CT with artificial intelligence could make detecting and quantifying these nematodes even faster and more accurate. This method could also be expanded to study other soil organisms, making it a valuable tool for monitoring soil health and improving agricultural pest management.

## Introduction

X-ray computed tomography (CT) has emerged as a promising tool in soil science due to its non-invasive, high-resolution, 3D imaging capabilities (Helliwell et al., 2013). This emerging method offers unprecedented insights into the complex network of soil pores (Tracy et al., 2012; Yang et al., 2018) in which soil-borne organisms may occupy and shape soil health and fertility.

Here, we demonstrate the potential for the deployment of X-ray CT to identify soil pathogens, focusing on plant-parasitic nematodes as a case study. Every major crop species is host to at least one plant-parasitic nematode species, resulting in >$470m in global crop losses per day in both developing and developed countries (Elling, 2013). *Globodera pallida* are obligate sedentary parasites of Solanaceous plants, particularly potato, and represent a large economic burden as a result of their highly specialised and efficient parasitism. Despite their impact, growers often underestimate their economic effect due to the lack of clear symptoms and the need for labour-intensive soil sampling. Extraction of cysts from the soil via flotation and elutriation can often lack efficiency and microscopic analysis of yielded cysts is laborious and time-consuming (Wainer & Dinh, 2021). Effective sampling requires soil samples from multiple points, making the process seasonally constrained (Been & Schomaker, 2000).

An alternative technique that is rapid, efficient, and accurate would optimise detection thereby improving advisory procedures. Reliable quantification of *G. pallida* populations directly influences control strategies, including crop rotation, nematicide application, and the selection of resistant varieties. Nematicide toxicity has led to the withdrawal of several products due to environmental concerns (Sasanelli et al., 2021) and placed increased strain on legislative measures, such as the European Council Directive and North American Plant Protection Organization (NAPPO) regulations, aiming to control *G. pallida* spread. The European and Mediterranean Plant Protection Organization (EPPO, 2017) has published methods for extracting *G. pallida* cysts, emphasising the importance of accurate quantification and viability assessment for effective management (Pillai & Danduran, 2019). Despite these efforts, *G. pallida* infestations continue to rise globally (He et al., 2022). Early detection and identification of pathogens are crucial to mitigate agricultural losses, necessitating a paradigm shift in detection methodologies.

Here, we apply this approach to identifying and quantifying *G. pallida* in different soil types, providing evidence for the capability of X-ray CT in soil pathogen detection and highlighting the potential of X-ray CT for identifying soil pests. Importantly, this method scans the entire sample rather than isolating cysts for analysis. With further optimisation, this approach could enable rapid soil analysis for multiple pathogens, helping to quantify pathogens and understand soil factors that affect pathogen communities.

## Results

### Cyst characterisation by X-ray CT

X-ray analysis enabled the detailed visualisation of *G. pallida* cysts, both externally and internally (Fig 1). Preliminary efforts determined that dry cysts possessed a dehydrated and impermeable cuticle, making it harder to discern the internal contents, resulting in an opaque core image of the cuticle (Fig 1A). Conversely, wet cysts exhibited a highly permeable and translucent cuticle, facilitating the precise observation of internal eggs (Fig 1B, C; S1 appendix). Cysts displayed a structurally sound outer cuticle and an egg-filled inner cavity with residual empty spaces, indicating a strong and intact structure.

**Fig 1.**
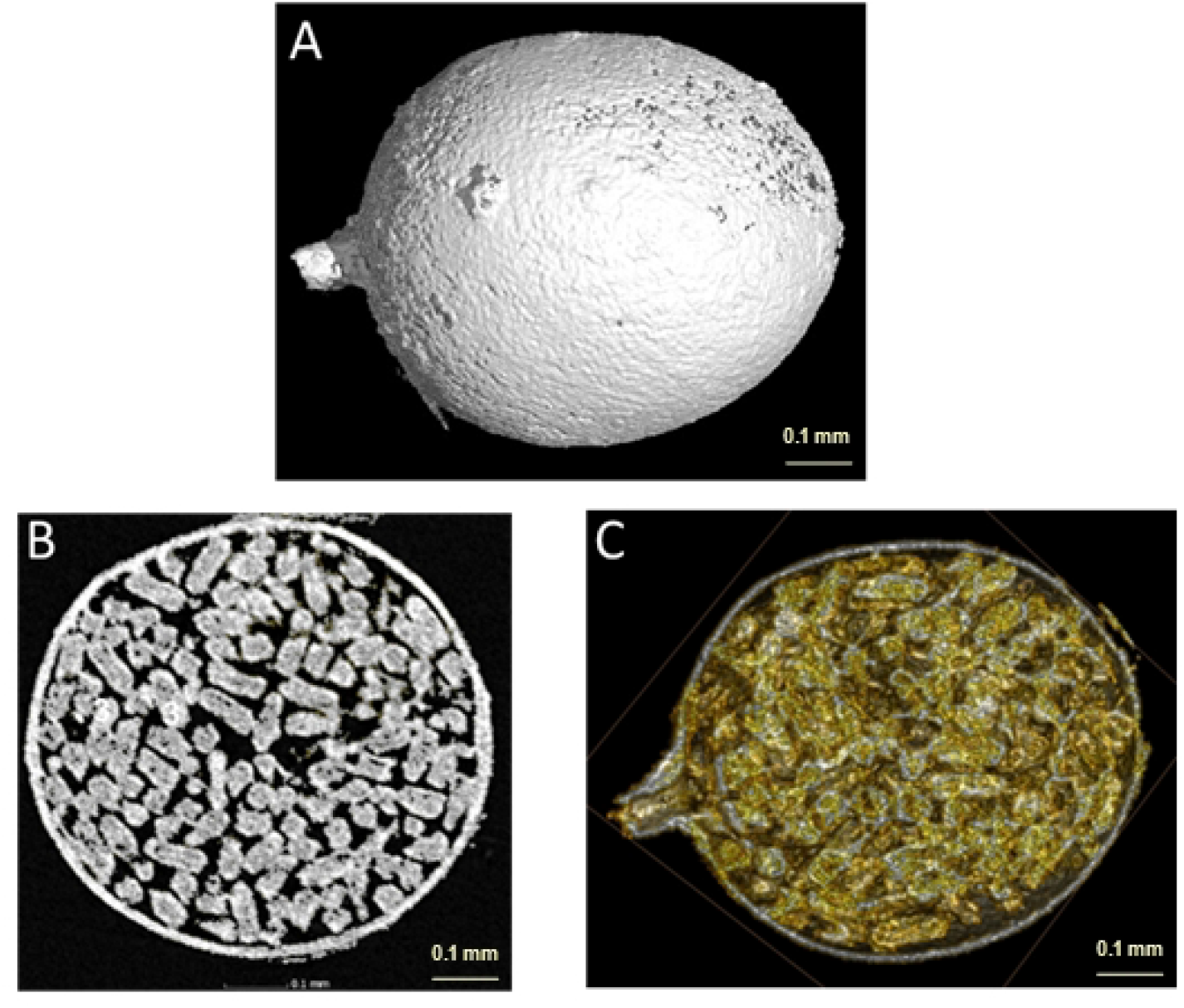
X-ray Computer Tomography scanning of a *Globodera pallida* cyst. *G. pallida* cysts were scanned via X-ray CT to visualise the cuticle (A) and interior contents from 360^0^ (B, C). Cross-sectional photos at 50% depth show distinct egg profiles (B, C).

### Identification of *G. pallida* in soil by X-ray CT

Several soil samples were spiked with varying unknown cyst quantities and analysed to evaluate the efficacy of X-ray CT methodology in identifying *G. pallida* cysts in soil. The presence of *G. pallida* cysts was confirmed in both sandy (Fig 2A, B) and loamy soil types (Fig 2C, D) despite inherent contrasts in soil composition, characterised by distinct textures and varying porosity levels. The detection of cysts proved straightforward, allowing for computer-led differentiation between damaged cysts and those that were intact (Fig 2E, S2 appendix).

**Fig 2.**
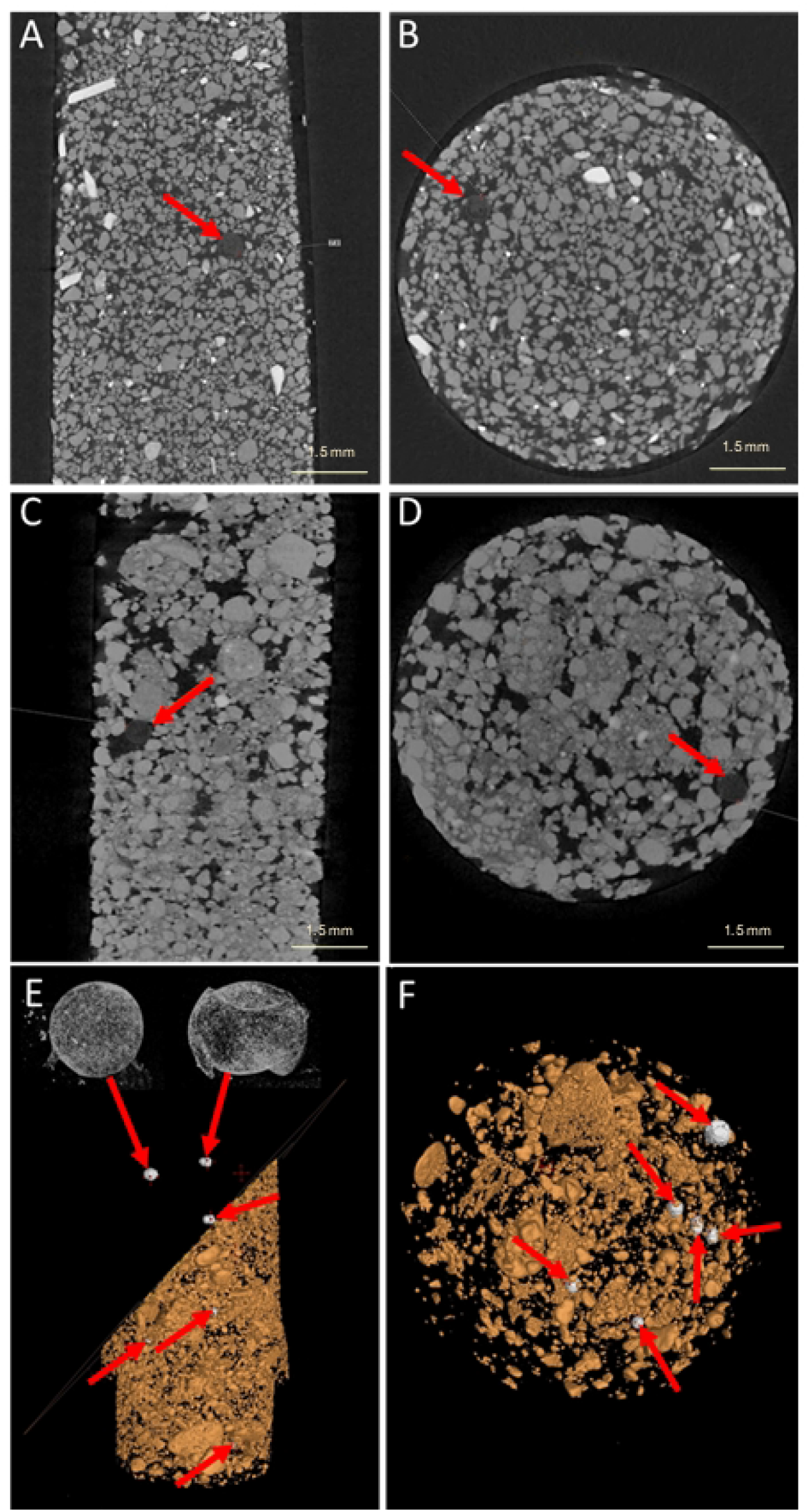
X-ray Computed Tomography scanning of soil containing *Globodera pallida* cysts. Sand (A, B) and loamy soil (C, D, E, F) were spiked with *G. pallida* cysts (red arrows) and scanned via X-ray CT. Soils were scanned in 15 ml Falcon tubes and visualised as either side (A, C, E) or top-down (B, D, F) profiles. Computer-led subtraction of soil (E) allowed distinct separation and quantification of cysts from the soil.

### Correlation between cyst volume and number of eggs

Previous work has determined a strong correlation between cyst-projected surface area and egg content (Urwin et al., 1997). X-ray CT allowed the quantification of cyst volume, ranging from 0.02 to 0.21 mm^3^. A positive correlation between cyst volume and egg count was observed (Pearson correlation analysis coefficient (R) value of 0.9882) (Fig 3). X-ray CT predictions of egg content demonstrated a +/-17% error margin compared to traditional egg counting methods.

**Fig 3.**
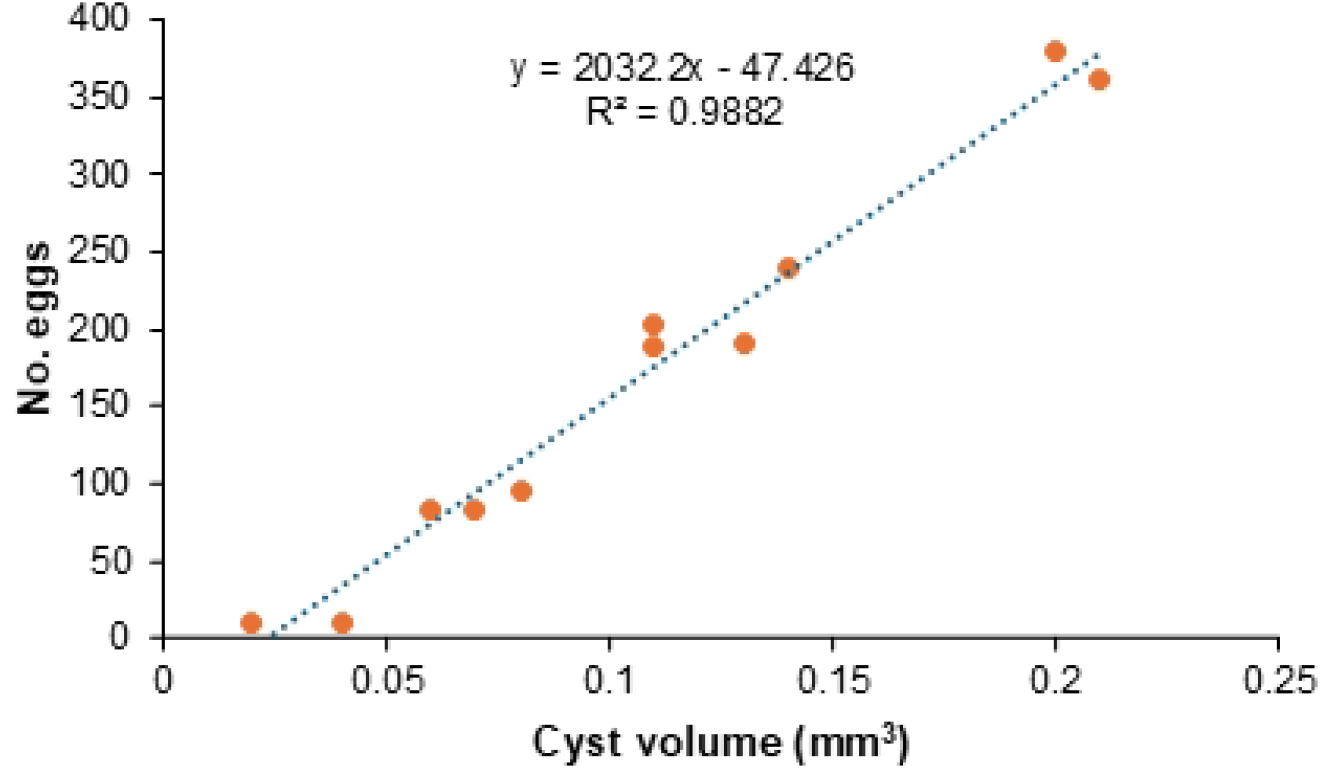
Egg number correlates with X-ray CT determined cyst volume. Cyst volumes were quantified through X-ray CT scanning, and exact egg numbers were determined via microscopy counts. Pearson correlation R^2^ = 0. 9882

## Discussion

This study provides the first insights into utilising X-ray Computed Tomography (CT) for identifying and analysing soil pathogens, with *G. pallida* as a case study. The primary advantage of this technique lies in its significant labour-saving potential compared to traditional cyst and egg extraction methods. Unlike conventional approaches, which require laborious sample handling and expertise (Ebrahimi et al., 2015; Reid et al., 2015; Christoforou et al., 2014), X-ray CT offers a rapid and straightforward procedure that does not demand specialised skills due to its potential to be computer-led. Operators can be easily trained to complete this task, making it a user-friendly tool in agricultural contexts. It also presages the easy detection of other important soil-borne organisms.

The process of using X-ray CT is not only efficient but also highly adaptable, making it suitable for various soil types and conditions. This adaptability enhances its potential for widespread use in agricultural monitoring. A key strength of X-ray CT is its ability to provide detailed three-dimensional imaging of *G. pallida* cysts, which significantly enhances our understanding of their morphology and internal structures. These insights are crucial for understanding the physiological states and viability of *G. pallida* (Price et al., 2021), aiding in developing effective pest management strategies.

X-ray CT extends the earlier work of Atkinson et al. (2001), who analysed cross-sectional areas and egg numbers by enabling the measurement of cyst volumes and providing more accurate egg counts. Our findings demonstrate that the number of eggs tends to increase proportionally as cyst volumes increase. This suggests that cysts are typically filled with eggs and are not larger than necessary to contain the observed number of eggs. Such detailed data are invaluable for advancing our knowledge of *G. pallida* biology, life cycles, and reproductive dynamics.

One of the most transformative aspects of X-ray CT is its potential for automation through artificial intelligence (AI) (Graas et al., 2023). AI can be trained to identify cyst images, further streamlining the process and reducing the need for manual intervention. This integration could accelerate the analysis of X-ray CT data and enable the automation of complex tasks. Machine learning algorithms can be developed to enhance the precision of pathogen detection, facilitating rapid and accurate identification of nematode populations.

Beyond its application to *G. pallida*, X-ray CT holds promise for concurrently detecting other nematodes and invertebrate pests and pathogens. Its versatility makes it a valuable tool for comprehensive soil health assessments, contributing to sustainable pest management in agriculture. We are currently investigating the utility of the technique in identifying other important soil-borne organisms. While some of these may be too small for direct visualisation, X-ray CT can indirectly detect their effects by visualising changes in root morphology, soil structure and disease symptoms caused by pathogenic activity (Hou et al., 2022; Teramoto et al., 2020).

However, the technology is not without limitations. While the resolution of X-ray CT is sufficient for detecting *G. pallida* cysts, further refinement may be necessary to improve the detection of smaller pathogens or early-stage nematodes. Future research should address this limitation by enhancing the sensitivity and resolution of X-ray CT technology.

## CONCLUSION

In conclusion, adopting X-ray CT for soil pathogen detection offers numerous advantages over existing methods, including the opportunity for holistic AI-led soil health analysis. Its application extends beyond *G. pallida* to other nematodes and invertebrate pests, making it a critical technology for improving our understanding and management of soil-borne threats in agricultural systems. By providing a non-invasive, high-resolution imaging solution, X-ray CT represents a significant advancement in the tools available to researchers and practitioners in nematology and soil science. Its success in detecting *G. pallida* underscores its potential for broader applications in identifying other soil-borne pathogens, reinforcing sustainable agricultural practices, and advancing soil management strategies.

## Material and methods

### Globodera pallida cyst samples

*G. pallida* (Lindley) cysts were spiked into sand (RHS Horticultural Sharp Sand) or loam-based (Baileys of Norfolk) soils in 15ml Falcon tubes. Cysts were handpicked from soils and then crushed in water to quantify their egg content via microscopy.

### X-ray Computer Tomography Scanning

*G. pallida* and soil samples in 15 mL falcon tubes were characterised using the GE Nanotom M X-ray CT machine (GE Measurement and Control Solutions, Germany). The v|tome|x M was set at a voltage of 65 kV and a current of 300 μA to optimise contrast between background soil and objects of interest. The ‘Fast Scan option’ achieved a voxel resolution of 1.60 μm. 1,078 projection images were taken per scan at 200 m/s per image. Once scanning was complete, the images were reconstructed using Phoenix datos|×2 rec reconstruction software, combining the scans into a single 3D volume representing the entire core.

### Image Processing

Image analysis for X-ray CT images was performed using the software VGStudioMax®, version 3.2 (Volume Graphics GmbH, Heidelberg, Germany) to segment the samples. The samples were segmented by setting seed points and using selected threshold values in the Region grower, enabling the selection of grey-scale pixels about cyst materials. Once the cysts were segmented from the image, the erosion and dilation tool was selected at 1 pixel using the Region Growing tool. A comprehensive image analysis protocol was developed, which included image segmentation, feature extraction, and *G. pallida* classification. *G. pallida* cysts were identified based on their distinctive morphology, including size, shape, and density characteristics observed in the X-ray CT images.

## Acknowledgement

We gratefully acknowledge funding from the Leverhulme Trust (RPG-2019-162).

## Supporting information

**S1 Appendix –** 3D X-ray Computer Tomography of *Globodera pallida* cyst.

**S2 Appendix –** 3D X-ray Computer Tomography of the distribution of *Globodera pallida* cysts in a soil sample.

## References

Atkinson HJ, Holz RA, Riga E, Main G, Oros R, Franco J. An algorithm for optimising rotational control of Globodera rostochiensis on potato crops in Bolivia. J Nematol. 2001 Jun;33(2-3):121–5.

Been TH, Schomaker CH. Development and evaluation of sampling methods for fields with infestation foci of potato cyst nematodes (Globodera rostochiensis and G. pallida). Phytopathology. 2000;90:647–56. Doi: doi: 10.1094/PHYTO.2000.90.6.647

Christoforou M, Pantelides IS, Kanetis L, Ioannou N, Tsaltas D. Rapid detection and quantification of viable potato cyst nematodes using qPCR in combination with propidium monoazide. Plant Pathol. 2014;63:1185–92. doi: 10.1111/ppa.12193.

Ebrahimi N, Viaene N, Moens M. Optimizing trehalose-based quantification of live eggs in potato cyst nematodes (Globodera rostochiensis and G. pallida). Plant Dis. 2015;99(5):667–72. doi: 10.1094/PDIS-09-14-0940-RE.

Elling, A. A. (2013). Major emerging problems with minor meloidogyne species. Phytopathology, 103(11), 1092–1102. 10.1094/PHYTO-01-13-0019-RVW

EPPO. Nematode extraction PM 7/119(1). European and Mediterranean Plant Protection Organization. EPPO Bull. 2013;43(3):471–95.

Graas A, Coban SB, Batenburg KJ, et al. Just-in-time deep learning for real-time X-ray computed tomography. Sci Rep. 2023;13:20070. doi: 10.1038/s41598-023-46028-9.

He Y, Wang R, Zhao H, et al. Predicting potential global distribution and risk regions for potato cyst nematodes (Globodera rostochiensis and Globodera pallida). Sci Rep. 2022;12:21843. doi: 10.1038/s41598-022-26443-0.

Helliwell JR, Sturrock CJ, Grayling KM, Tracy SR, Flavel RJ, Young IM, et al. Applications of X-ray computed tomography for examining biophysical interactions and structural development in soil systems: A review. Eur J Soil Sci. 2013;64(3):279–97. doi: 10.1111/ejss.12028.

Hou L, Gao W, der Bom F, Weng Z, Doolette CL, Maksimenko A, et al. Use of X-ray tomography for examining root architecture in soils. Geoderma. 2022;405:115405. doi: 10.1016/j.geoderma.2021.115405.

OEPP/EPPO. EPPO data sheets on quarantine pests [Internet]. 2014 [cited Year Month Day]. Available from: http://www.eppo.int/QUARANTINE/nematodes/Globodera_pallida/HETDSP_ds.pdf?utm_source=www.eppo.org&utm_medium=int_redirect.

Pillai SS, Dandurand LM. Evaluation of fluorescent stains for viability assessment of the potato cyst nematodes Globodera pallida and G. ellingtonae. Adv Biosci Biotechnol. 2019;10:244–58. doi: 10.4236/abb.2019.108019.

Price JA, Ali MF, Major LL, Smith TK, Jones JT. An eggshell-localised annexin plays a key role in the coordination of the life cycle of a plant-parasitic nematode with its host. PLoS Pathog. 2023;19(2). doi: 10.1371/journal.ppat.1011147.

Sasanelli N, Konrat A, Migunova V, Toderas I, Iurcu-Straistaru E, Rusu S, et al. Review on control methods against plant parasitic nematodes applied in southern member states (C zone) of the European Union. Agriculture. 2021;11:602. doi: 10.3390/agriculture11070602.

Reid A, Evans F, Mulholland V, Cole Y, Pickup J. High-throughput diagnosis of potato cyst nematodes in soil samples. Methods Mol Biol. 2015;1302:137–48. doi: 10.1007/978-1-4939-2620-6_11.

Tracy SR, Black CR, Roberts JA, Sturrock C, Mairhofer S, Craigon J, et al. Quantifying the impact of soil compaction on root system architecture in tomato (Solanum lycopersicum) by X-ray micro-computed tomography. Ann Bot. 2012;110(2):511–9. doi: 10.1093/aob/mcs031.

Teramoto S, Takayasu S, Kitomi Y, et al. High-throughput three-dimensional visualization of root system architecture of rice using X-ray computed tomography. Plant Methods. 2020;16:66. doi: 10.1186/s13007-020-00612-6.

Urwin PE, Lilley CJ, McPherson MJ, Atkinson HJ. Resistance to both cyst and root-knot nematodes conferred by transgenic Arabidopsis expressing a modified plant cystatin. Plant J. 1997;12(2):455–61. doi: 10.1046/j.1365-313X.1997.12020455.x.

Wainer J, Dinh Q. Taxonomy, morphological and molecular identification of the potato cyst nematodes, Globodera pallida and G. rostochiensis. Plants. 2021;10(1):184. doi: 10.3390/plants10010184.

Yang Y, Wu J, Zhao S, et al. Assessment of the responses of soil pore properties to combined soil structure amendments using X-ray computed tomography. Sci Rep. 2018;8:695. doi: 10.1038/s41598-017-18997-1.

